# Capacitive coupling increases the accuracy of cell-specific tumour ablation by electric fields

**DOI:** 10.1101/819375

**Authors:** Terje Wimberger, Verena K. Köhler, Eva K. Ehmoser, Klemens J. Wassermann

## Abstract

Irreversible electroporation holds great potential for cell-specific lysis due to the size-dependent susceptibility of cells to externally imposed electric fields. Previous attempts at selective cell lysis lead to significant overlap between affected populations and struggle with inconsistent biological outcome. We propose that charge transfer at the electrode-liquid interface is responsible by inducing multifactorial effects originating from both the electric field and electrochemical reactions. A promising remedy is the coating of electrodes with a high-k dielectric layer. The resulting capacitive coupling restores the selective potential of electric field mediated lysis in a microfluidic setup. Initial experiments show the consistent depletion of erythrocytes from whole blood while leaving leukocytes intact. The same is true for the reproducible and selective depletion of Jurkat and MCF-7 cells in a mixture with leukocytes. Unexpectedly, the observed order of lysis cannot be correlated with cell size. This implies that the cellular response to capacitive coupling features a selective characteristic that is different from conventional lysis configurations.

## 1. Introduction

Biological membranes exposed to an electric field of sufficient magnitude undergo electroporation, corresponding to an increase in permeability of the lipid bilayer [1,2]. If the field is further increased and the transmembrane potential reaches a critical threshold, the effect becomes irreversible and leads to cell lysis [3,4]. The known factors governing changes in the local transmembrane potential are described by

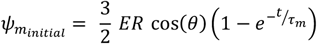

where *E* is the externally applied electric field, *R* is the cell radius, *θ* being the angle between the electric field direction and the point on the membrane affected by the field, *t* being the field exposure time and *τ*_*m*_ being the membrane charging time [5,6]. Since the change in transmembrane potential is co-determined by cell size, a window of opportunity for selective cell lysis is implied and has been exploited for selective lysis in previous occasions [7,8].

To our knowledge, three previous studies have attempted the electric field-mediated lysis of malignant cells from blood. The first study featured in *Nature Biotechnology* was confined to a batch configuration [8]. CMK tumour culture cells were spiked with peripheral blood mononuclear cells and showed a higher susceptibility to electric fields. At settings that deplete > 98% of CMK cells, monocyte viability is reduced below 20%. Shortcomings in the electrical setup result in 67% confidence intervals for 7 repetitions and the need for a water-cooling system to counter significant joule heating. The second study was featured in *Blood* and shows the efficient depletion of multiple myeloma cell lines from blood [9]. While it was shown that stem cell function of the surviving population was preserved, less than 1% of monocytes and 10% of lymphocytes survived the treatment. In the third study published in *Lab on a Chip*, 98% of M109 circulating tumour cells are eliminated at field magnitudes accompanied by loss of viability of > 50% of white blood cells [10].

The biggest obstacle for using electric fields in a cell-specific manner is the multifactorial nature of the effect. While the field effect might be considered specific, it is inevitably accompanied by a number of non-specific side effects caused by electrochemical events along the electrode-suspension interface [11]. These include pH- and temperature changes as well as ionic radicals and gas formation, which in turn negatively affect reproducibility, cellular integrity and electrode lifetime [12,13].

Strategies to minimize the impact of electrochemistry have included spatial separation of electrodes [12]. A downside of large electrode distance is the high voltage required to reach a sufficient local electric field. Nanosecond pulse electroporation represents another method of tackling the issue of electrochemistry, but similarly requires high voltages and specialized equipment [14]. Microfluidic approaches to reduce the adverse effects of electrolysis encounter a different set of obstacles; complex geometries lead to loss of field uniformity and highly miniaturized systems are accompanied by cell deformation and limited throughput [15,16]. In summary, there is no consensus on the method of choice for applying uniform electric field effects to a sufficient number and fraction of cells.

Since all the discussed side effects are detrimental to a fraction of any given cell population, discussing the issue of cell specific lysis is somewhat misleading because the real challenge is posed by specifically keeping a certain cell population intact. In a previous publication, we presented an alternative approach to deal with unspecific side effects posed by electrochemistry; by coating electrodes with a high-k passivation layer, we were able to deliver sufficient electric fields to a cell suspension while significantly reducing electrochemical reactions along the interface [17]. This is a result of capacitive coupling which has a history of use in contactless dielectrophoresis. Here it has successfully resolved issues associated with electrode fouling during continuous low-energy field application at high frequencies [18,19].

The theoretical possibility of size discrimination was successfully put into practice for large differences in size, such as between bacteria and mammalian cells [17]. Subsequent work revealed that our device can specifically target cell populations of similar size. Sticking to our established model of whole blood (WB) lysis, we present the selective, field-mediated lysis of erythrocytes as a novel method for leukocyte isolation.

## 2. Materials and methods

### 2.1 Electrode passivation and prototype assembly

Details and optimization of the passivation procedure and design considerations are provided in a previous publication [17]. Briefly, commercial titanium foils (grade 2, cpTi, 99.2% purity, 110 μm thick) were cut and 0.8 mm holes drilled to form in- and outlet ports. Foils were cleaned in ultrasonic bath with a series of acetone, ultrapure water and isopropanol for 20 min each. This was followed by thermal oxidation in a muffle furnace (L 9/11 P330, Schaefer & Lehmann, Germany) for 3 hours at 650 °C with 2 hours rise-time and passive cooling to RT. Microfluidic cell lysis units were assembled by cutting the flow chamber geometry (Scheme 1) into double-sided 81.3 μm thick adhesive tape (Arcare 90445, Adhesive Research, Ireland), sandwiched by two pieces of passivated titanium foil. The adhesive features continuous thickness, bio-compatibility [20] and yields a channel volume of 10 μl. Adaptors to the flow system (NanoPort Std 6-32 Coned 1/32, IDEX, USA) were fixed to in- and outlets. Circuit closure was ensured by removal of outer oxide layer from the bottom sides of both electrodes using diamond files until the resistance dropped below 1 Ohm. Every new prototype was assessed in term of fluidics and electrical performance by application of standard pulses when filled with PBS.

**Scheme 1:**
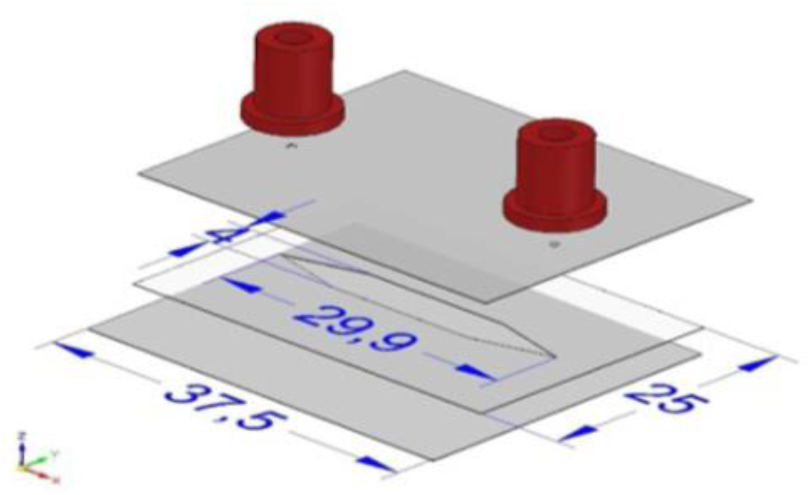
Geometric design of the electric cell lysis unit. The fluidic channel is formed by double-sided adhesive tape. All measurements in mm.

### 2.2 Preparation of working solutions

Preparation of the electroporation buffer (EPB) was performed by incremental addition of PBS to autoclaved 250 mM sucrose solution until conductivity reached 100 μS/cm. WB samples were directly diluted in 250 mM sucrose without conductivity manipulation. Conductivity was measured using a conductivity meter (B-771 LAQUAtwin, HORIBA Advanced Techno, Japan). WB was collected from healthy volunteers using K3-EDTA collection tubes (Vacuette, Greiner Bio One, Austria), immediately stored a 4 °C and kept for a maximum of 3 days. Any dilutions were prepared with 250 mM sucrose solution. Erythrocyte lysis buffer (ELB) containing 155 mM NH_4_Cl, 10 mM KHCO_3_ and 0.1 mM EDTA was prepared, sterilized by filtration (0.22 μm PVDF filter) and stored at 4 °C until use.

### 2.3 Cell culture

Jurkat T lymphocytes as a model for leukaemia (Clone E61, ATCC®TIB152™) were cultivated at 37 °C and 5% CO_2_ in RPMI (Thermo Fisher, 21875091) supplemented with 10% FBS (Thermo Fisher, 10500) and 1% Pen/Strep Antimycotic-Antimycotic (Thermo Fisher, 15240). Jurkat T lymphocytes suspension cultures were passaged by transferring a fraction of the suspension to a 15 ml tube, centrifuging for 5 min at 400 rcf (RT, Eppendorf 5430; Rotor: F-35-6-30), re-suspension in fresh medium and transfer to cultivation flask.

MCF-7 cells as a model for circulating tumour cells (Public Health England, 86012803, Lot No. 14I018) were cultivated at 37 °C and 5% CO_2_ in MEM (Thermo Fisher, 21875091) supplemented with 10% FBS (Thermo Fisher, 10500), 2% L-Glutamine (Thermo Fisher, 25030081), 1% non-essential amino acids (Thermo Fisher, 11140050) and 1% Pen/Strep Antimycotic-Antimycotic (Thermo Fisher, 15240). MCF-7 were passaged by washing with PBS (1x from stock: Thermo Fisher, 70011044) followed by trypsinization (0.25%, Thermo Fisher, 25200) for 5 min at 37 °C. Any sterile protocols were processed in biological safety cabinets (Herasafe KS, Class II, Thermo Fisher, 51022488).

### 2.4 Leukocyte isolation

10 ml ELB was mixed with 1 ml of WB, incubated 10 min at RT and inverted repeatedly. The suspension was centrifuged at 500 rcf for 10 min RT (Eppendorf 5430; Rotor: F-35-6-30). These steps were repeated until a white cell pellet was obtained (indicating erythrocyte depletion). After washing twice with sucrose-PBS solution set to 100 μS/cm, cells were counted, and concentration was adjusted to 5×10^5^ cells/ml. Viability was assessed by staining with Hoechst 33342 (Ref.: B2261, Sigma-Aldrich Co. LLC.). Preparations with less than 80% viability were discarded.

### 2.5 Preparation of Jurkat T lymphocytes and MCF-7 culture cells

Jurkat T lymphocyte suspension was centrifuged at 400 rcf for 5 min and excess medium discarded. Cells were re-suspended in 5 ml EPB (100 μS/cm). This step was repeated three times. During the last centrifugation step, an aliquot of cells was counted and re-suspended in the amount of EPB needed for a final cell concentration of 5×10^5^ cells/ml and the suspension conductivity recorded. Adherent MCF-7 cells were washed with PBS, trypsinized for 5 min at 37 °C and re-suspended in culture medium. Further preparation steps were performed analogous to Jurkat preparation.

### 2.6 Preparation of spike-in suspensions

For the WB-leukocyte spike-in experiments, leukocytes prepared as in section 2.4 were equilibrated in EPB with a conductivity of 210 μS/cm and a final concentration of 10^6^ cells/ml. WB was diluted 1:500 in EPB (210 μS/cm) and counted. Further EPB was added to adjust cell concentration to 10^6^ cells/ml. Both suspensions were mixed 1:1 prior to field exposure. For the Jurkat T lymphocyte-spiked leukocyte suspension, both populations were prepared as described in sections 2.4 and 2.5, but with an increased concentration of 10^6^ cells/ml, respectively. Jurkat-T lymphocytes in EPB were stained with 10 nM Calcein-AM solution (Thermo Fischer, C3100MP) for 60 min at 37 °C to allow for population tracking in flow cytometry. An aliquot of unstained cells was set aside for flow cytometry negative controls. Prior to pulse application, stained Jurkat T lymphocytes were mixed 1:1 with leukocytes for final concentrations of 5×10^5^ cells/ml each. For the MCF-7-spiked leukocyte suspension, both populations were prepared as described in sections 2.4 and 2.5 with cell counts increased to 10^6^ cells/ml. MCF-7 cells in EPB were stained with a 1:20 dilution of FITC Anti-human CD326 (Biolegend, Cat.no. 324204) for 30 min at 4 °C to allow for population tracking in flow cytometry. An aliquot of unstained cells was set aside for flow cytometry negative controls. Prior to pulse application, stained MCF-7 are mixed 1:1 with leukocytes for final concentrations of 5×10^5^ cells/ml each. Live-dead discrimination of the MCF-7 population was assessed by Hoechst 33342 staining (1 μg/ml) prior to data acquisition.

### 2.7 Electric field application

1 ml of respective cell suspensions were transferred to a 1 ml syringe (Omnifix-F, Braun, Germany) and injected to the electric cell lysis unit (ECLU) by a syringe pump (Fusion 200 Touch, KR Analytical Ltd, United Kingdom) set to a flow rate of 100 μl/min. To discriminate between parameters, at least five ECLU chamber volumes - a total of 50 μl - were passed through the device after any parameter change and before collecting an aliquot for further analysis. Given the channel volume of 10 μl, a flow rate of 100 μl/min means that cells are exposed to the indicated field parameters for 6 seconds. A frequency of 100 Hz results in 600 square wave periods. Respective voltages refer to square waves applied by a function generator (DG4102, Rigol) connected to a voltage amplifier (Falco WMA-300, Falco Systems, Netherlands). Voltage and current (via a 2 Ω resistor) were monitored by an oscilloscope (DS1104B, Rigol).

### 2.8 Data acquisition and analysis

Population viability was assessed by transferring a 10 μl aliquot of ECLU-treated cell suspensions to a haemocytometer (Thoma, Optik Labor) and imaging with a digital camera (Prosilica GT, Allied Vision) mounted on an inverted microscope (CKX41 Fluo V2, Olympus). Bright-field images were recorded for total cell count. For leukocytes, Jurkat T lymphocytes and MCF-7 cells, 1 μg/ml Hoechst 33342 viability dye was added to discriminate dead cells. For viability assessment of leukocytes in WB-leukocyte spike-in experiments, 1 μg/ml of membrane impermeable Propidium Iodide (PI) was added after field exposure and prior to imaging. Flow cytometry data acquisition of (Jurkat T lymphocyte and MCF-7) spike-in experiments was performed with BD FACSCanto II. Forward scatter (FSC) and side scatter (SSC) thresholds were set to eliminate cell debris from the final readout. 10 000 events were recorded for each parameter. Ca-AM and CD326-FITC parameters were recorded in the FITC channel, Hoechst 33342 staining was recorded in the Pacific Blue channel. Data was gated in Flowing Software 2.5.121 and statistical evaluation and data visualization carried out with Graphpad Prism 7. Where applicable, standard deviation (SD) refers to standard deviation of all replicates and n refers to technical replicates. Flow cytometry data was composed from biological replicates as indicated.

## 3. RESULTS AND DISCUSSION

### 3.1 High-k passivation limits joule heating

A persisting issue in microfluidic, electric field applications is the limited throughput of batch configurations and the complexity of circumventing negative side effects of electrochemistry in flow-through setups [16]. Fig. 1 shows the temperature stability of the flow-through design (Scheme 1) with and without high-k passivated electrodes for 10 min of continuously applied square wave pulses (30 V at 100 Hz). Without passivation, data shows a temperature increase of 16.3 °K for processing 1 ml of sample. A 10 °K increase is reached within 2 minutes of field exposure, despite the moderate cooling effect of the sample injection. The highest sensor noise recorded was Δ0.16 °K. With high-k passivation, a maximum temperature increase of 0.67 °K after 540 seconds of field application is observed. This value accounts for the processing of 1 ml of sample at 100 μl/min at a voltage sufficient for all selective lysis applications in the following series of experiments. The negligible temperature increase upon field application showcases the efficiency of high-k passivation in terms of energy conservation.

**Fig. 1.**
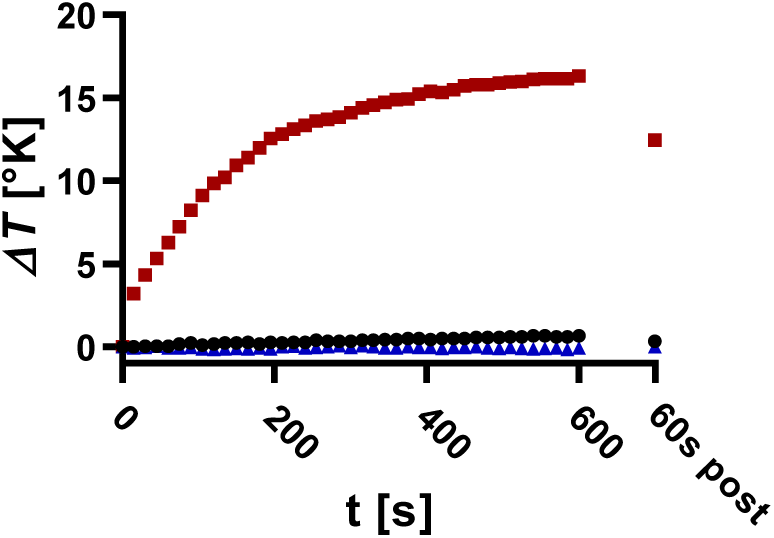
Temperature drift from starting temperature with 30 V applied continuously at 100 Hz with (·) and without passivation (▪) compared to thermometer drift with no field applied (▴).

### 3.2 Electric field mediated erythrocyte lysis is conductivity dependent

This section shows erythrocyte lysis at varying field strength and buffer conductivity as a result of WB dilution with 250 mM sucrose. Previous examinations of the correlation between buffer conductivity and electric field mediated lysis have resulted in contradicting reports; Pucihar *et al.* reported reduced field effects on DC3F Chinese hamster fibroblasts when reducing medium conductivity in the range of 0.01 - 16 mS/cm [21]. Such negative correlations are further found by *Ivorra et.al.* [22] and *Rols and Teissié* [4]. Contradictingly, *Djuzenova et al.* found that reducing conductivity between 0.8 - 14 mS/cm increased electroporation of murine myeloma cell line Sp2/0-Agl4 [23]. Such a positive correlation was further found by *Müller et. al.* [24] and *Silve et. al.* [25]. These studies do not cover the lower conductivity range of 108 - 5300 μS/cm tested here.

Results in fig. 2 indicate a clear correlation between erythrocyte lysis and solution conductivity. It is worth noting that SD negatively correlates with conductivity, as is described in literature for much larger changes in buffer composition [21]. Since dilution also reduces cell concentration, a highly conductive 1:100 dilution with PBS confirms the difference in lysis is due to the conductivity change and not the cell density. This in turn makes lysis of high-density samples feasible if their conductivity can be sufficiently lowered to allow for electric field penetration.

**Fig. 2.**
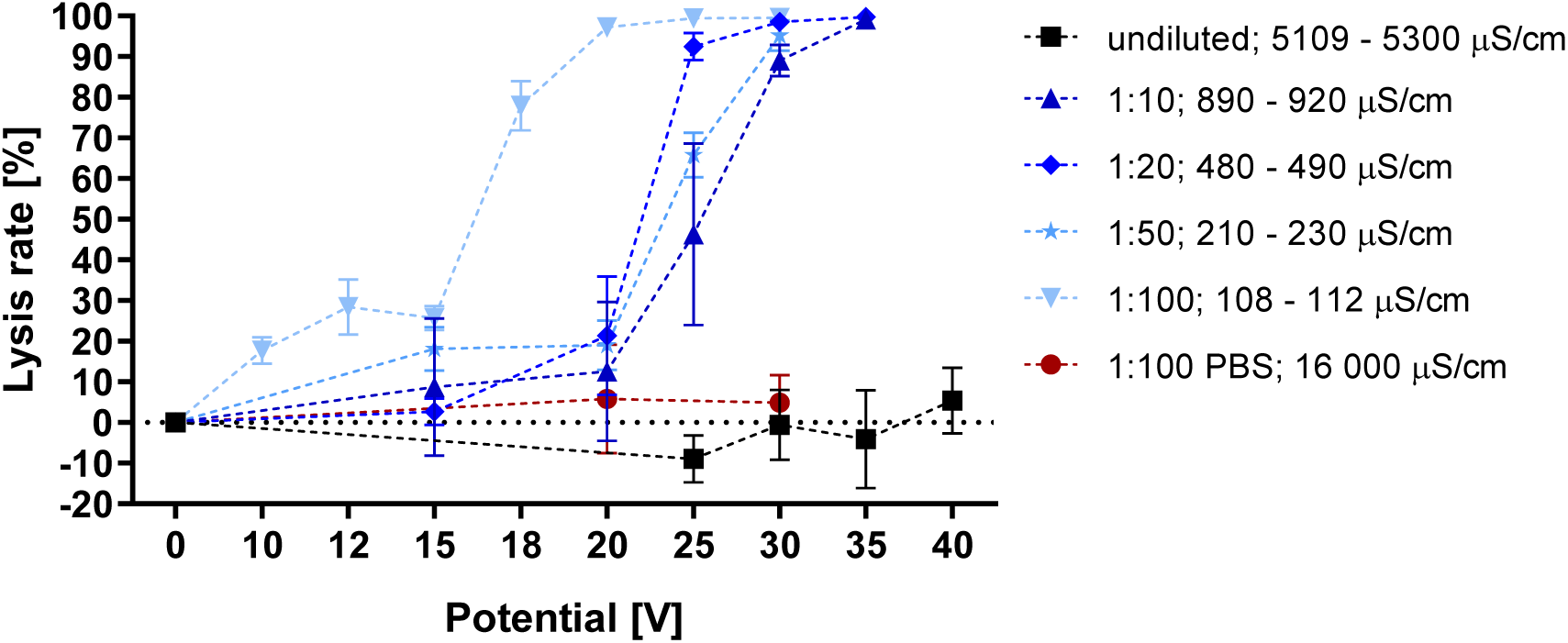
Voltage dependent lysis rate of various WB dilutions from the same donor. Conductivity range refers to start and end of the experiment. λ: 100 Hz; flow rate: 100 μl/min; n = 3

### 3.3 Leukocyte membrane integrity is preserved during erythrocyte lysis

We proceed to investigate the lysis parameters of WB and leukocyte populations separately prepared from WB (Fig. 3a). A field effect acting on erythrocytes can be observed at 18 V (77.9% lysis) and complete erythrocyte lysis is achieved at 25 V (99.4% lysis). Further increasing the potential to 30 V does not significantly impact lysis of remaining cells (99.5% lysis) because WB also contains a residual fraction of leukocytes. With rising field strength, lysis of the isolated leukocyte population decreases to 77.1% at 40 V. These results reveal a wide window of opportunity for selective enrichment of leukocytes. It is curious that the lysis curve for leukocytes spans over 20 V potential difference or twice the magnitude of lysis onset (Fig. 3a). Other studies find that leukocyte lysis occurs in a narrow window of 1/4^th^ increase in electric field strength from onset of lysis. In a specific instance this range lies between 0.4 – 0.5 kV/cm [10].

**Fig. 3:**
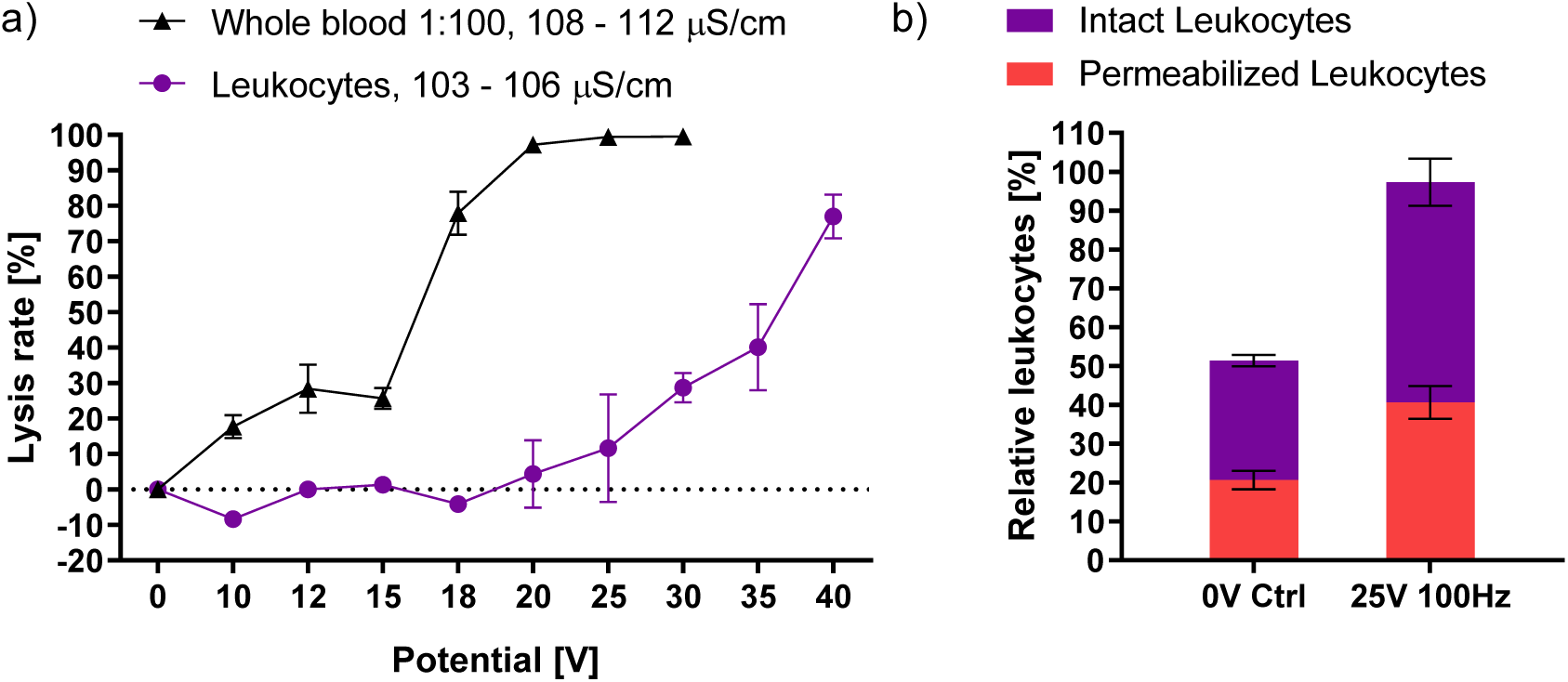
(a) Voltage-dependent lysis rate of isolated leukocytes in EPB. Erythrocyte lysis at the same conductivity is taken from fig. 2 for comparison. 5×l0^5^ leukocytes/ml; λ: 100 Hz; flow rate: 100 nl/min; n = 3. (b) Enrichment of leukocytes in a leukocyte spiked WB dilution. Intact: Hoechst 33342 positive; Permeabilized: Propidium Iodide positive; Cell concentrations: 5×10^5^ cells/ml; σ: 220 - 230 μS/cm; n = 3

These combined findings are in contrast to previous reports on electric-field mediated lysis of WB. Leukocyte lysis typically occurs at a far lower energy threshold than red blood cells, which require up to three times the electric field strength before irreversible electroporation leads to efficient rupture [10]. While previous studies conform to the theoretical background described by equation [1], the presented data suggests an inverse relationship between cell size and lysis threshold.

To further investigate this discrepancy, we proceeded to determine the membrane integrity of cells that appear unaffected in lysis experiments. For this purpose, a spike-in suspension of WB and isolated leukocytes was prepared with a final cell ratio of 1:1. Controls contained 51.4% leukocytes and 48.6% erythrocytes. Conductivity was adjusted to 220 μS/cm. and voltage set to 25 V with all other parameters unchanged. These settings eliminated 94.4% of erythrocytes from the elution, which was comprised of 97.3% remaining leukocytes (Fig. 3b). In the control sample not subjected to an electric field, 40.3% of the leukocytes are PI positive (20.7% of total), indicating membrane aberrations not caused by the electric field. After field application and selective erythrocyte lysis, the fraction of permeabilized leukocytes remains equal (41.7%). This indicates that the applied field has a negligible impact on membrane integrity of leukocytes and serves to support the curious observations from fig. 3b. These results imply the feasibility of using electric fields toward discrimination of erythrocytes and, consequently, the selective isolation of viable leukocytes.

### 3.4 Electric field mediated lysis of Jurkat T lymphocytes, MCF-7 and leukocytes is not size dependent when using capacitive coupling

In pertinent literature, the lysis of leukaemia- and circulating tumour cells occurs at thresholds lower than that of the leukocyte population [8,10]. Similarly, fig. 4a shows that Jurkat T lymphocytes have the highest susceptibility to voltage dependent lysis. Square wave pulses of 15 V result in lysis of 77.4% of Jurkat- and 69.3% of MCF-7 cells. Notably, this means that both cancer models display a higher field susceptibility than erythrocytes at the same field strength and conductivity (Fig. 2, 3a, 25.7% lysis). Leukocytes are mainly comprised of neutrophils (7.3 – 9.7 μm), lymphocytes (5.89 – 6.09 μm) and monocytes (7.72 – 9.99 μm) [26]. The diameter of MCF-7 cells ranges from 15 – 17 μm [27] and Jurkat T lymphocytes range from 10 - 13 μm [28]. While erythrocytes are significantly smaller (7.81 ± 0.63 μm) [29] their non-spherical shape could change the dynamics of their susceptibility to electric fields. If cell diameter were the primary factor governing electric field mediated lysis, we would expect to find Jurkat T lymphocyte lysis rate to fall somewhere between the two other examined cell types. Fig. 4b plots the size of tested cell populations against the voltage needed for disruption of 50% of the respective cell type (EV50). Jurkat T lymphocytes show the highest susceptibility to voltage dependent lysis (13.3 V), followed by MCF-7 (13.6 V), erythrocytes (16.9 V) and leukocytes (36.5 V). Fig. 4b shows that there is no significant correlation between lysis rate and cell size in this capacitively coupled configuration (R^2^ = 0.249).

**Fig. 4.**
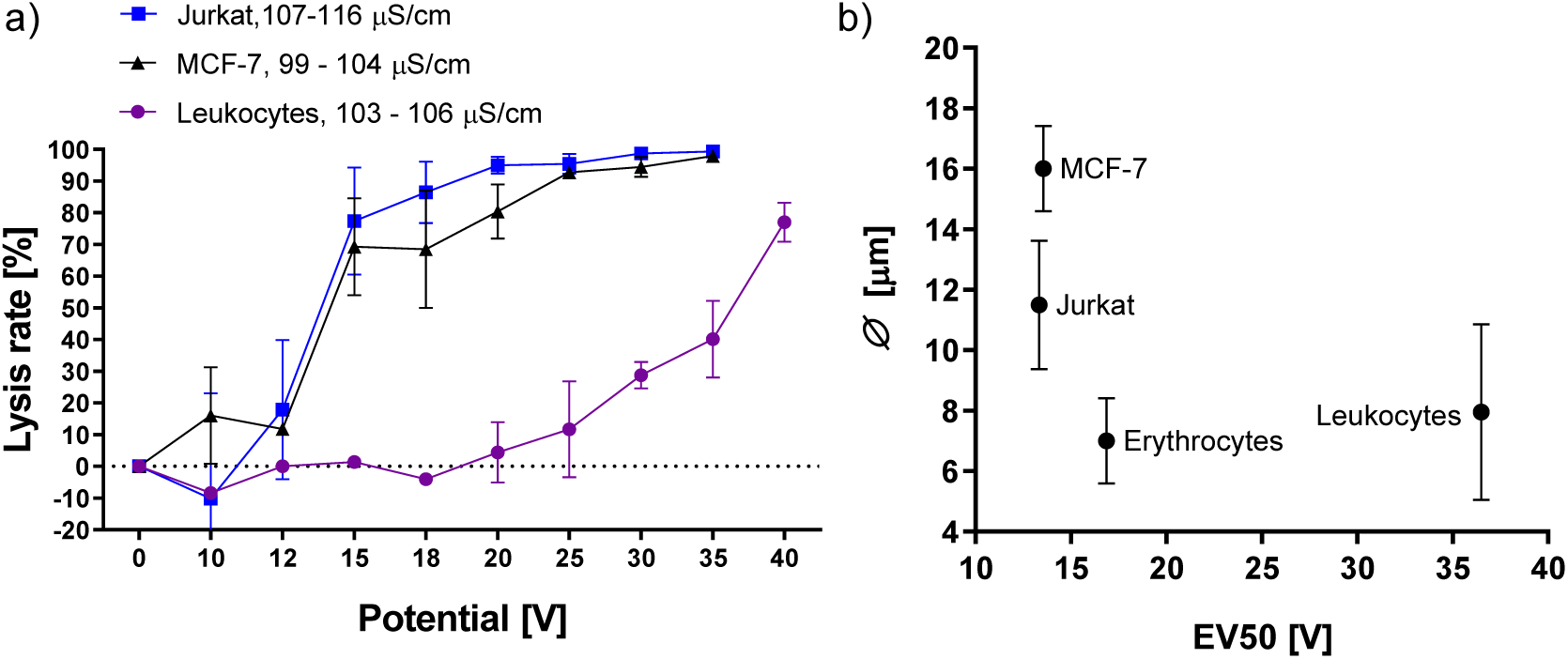
(a) Lysis rates of Jurkat T lymphocytes and MCF-7 cells exposed to increasing electric fields. Leukocyte lysis rate added from fig. 3a for comparison. Conductivity range refers to start and end of experiment. All cell concentrations: 5×10^5^ cells/ml; λ 100 Hz; flow rate: 100 μ/min; n = 3. (b) Average cell size according to literature plotted against effective voltage 50 (EV50), which is the voltage required for lysis of 50% of cells as determined by nonlinear curve fit (Boltzmann-sigmoid).

### 3.5 Capacitive coupling allows for selective elimination of Jurkat T lymphocytes and MCF-7 cells from a mixture with leukocytes

To test whether these effects persist in a mixed-population suspension, spike-in experiments were performed. Fig. 5 shows flow cytometry data with and without field application to a 1:1 mixture of Jurkat T lymphocyte- and leukocyte suspensions. In the mixed population without field application, 31.6% of counted events are Calcein-AM positive Jurkat T lymphocytes and 41.7% are identified as leukocytes by their SSC profile (Fig. 5a). Upon field application, < 0.1% of the events remain Calcein-AM positive Jurkat T lymphocytes while 50.7% of events are accounted to leukocytes (Fig. 5b). Cellular debris increases from 11.4% to 26.5% with field application (Fig. 5b,c). SSC gating and previous lysis experiments (Fig. 4a) support the notion that Jurkat T lymphocytes undergo lysis instead of losing Calcein-AM fluorescence. Fig. 5c shows combined event statistics from multiple experiments. The number of events attributed to unaffected Jurkat cells was below 0.1% in all three repetitions.

**Fig. 5.**
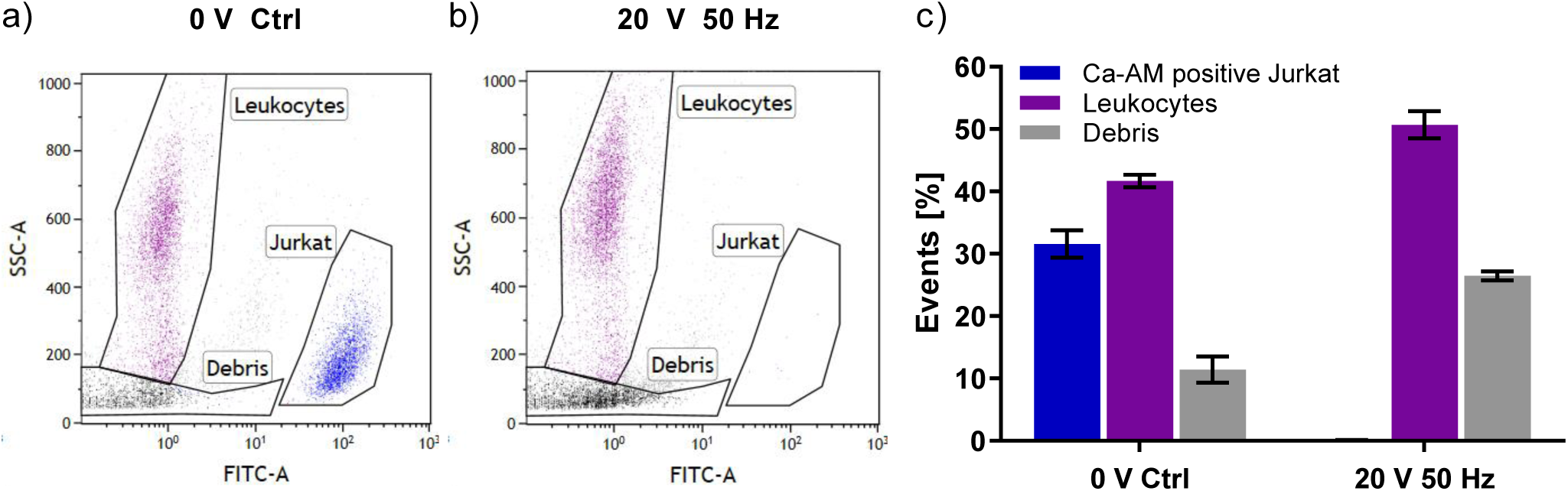
Selective elimination of Jurkat T lymphocyte population mixed with isolated leukocytes. Jurkat T lymphocytes were incubated with Calcein-AM prior to data acquisition and have high FITC fluorescence, (a) Mixed cell population without electric field application, (b) Mixed cell population after application of 20 V square wave pulses at 50 Hz. (c) Event statistics showing percentage of respective events. Conductivity: 97 μS/cm; events: 10 000; data composed from three biological replicates

Fig. 6a,b show flow cytometry data with and without field application to a 1:1 mixture of MCF-7 and leukocytes. MCF-7 cells were stained with FITC-conjugated antibody for identification. Lysis rate was assessed via Hoechst 33342 addition prior to data acquisition. In the mixed population control, 69.8% of labelled MCF-7 cells were counted as viable with 91.7% of leukocytes remaining intact (Fig. 6a,c). Application of 30 V square wave pulses at 100 Hz result in 97.9% lysis of the MCF-7 population while 68.5% of leukocytes remain intact (Fig. 6b,c). Fig. 6c shows the average population counts for two technical replicates with SD.

**Fig. 6.**
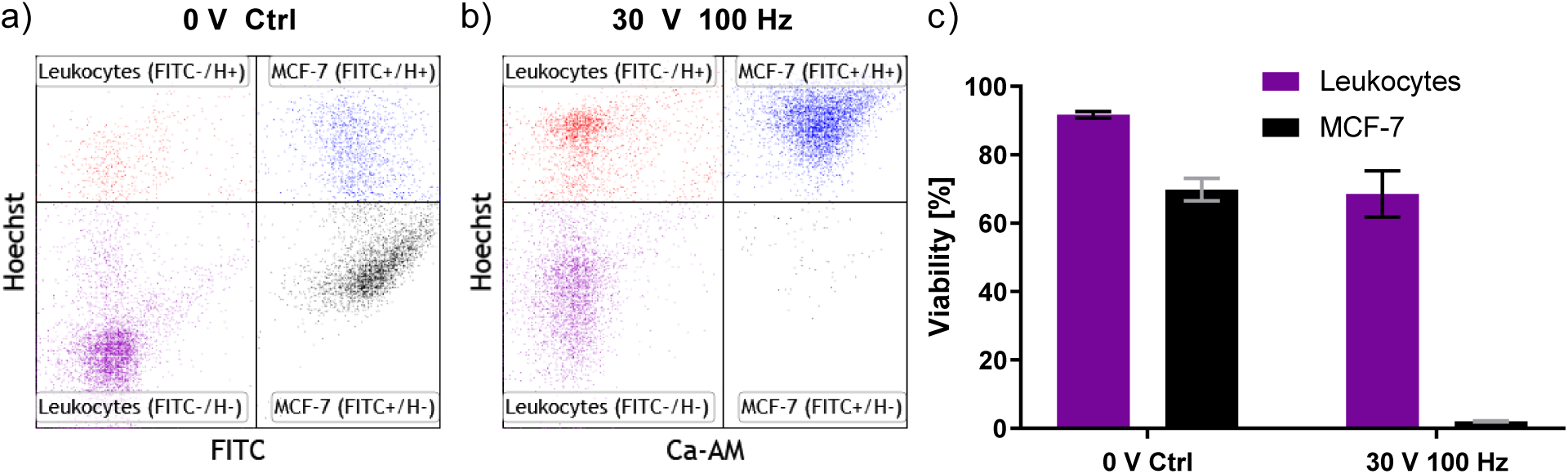
Selective elimination of MCF-7 cells mixed with leukocyte suspension, (a) Mixed cell populations without electric field, (b) Mixed cell suspension after application of 30 V at 100 Hz. (c) Statistics on viability after field application. Conductivity: 100-103 μS/cm; events: 10 000; data composed from two biological replicates

High-k dielectric passivation reliably reduces tumour cell count while keeping leukocyte populations intact. In the case of Jurkat T lymphocyte spike-in experiments, the leukocyte count remains virtually unchanged by the applied electric field. The higher energy pulses of 30 V applied to the MCF-7 leukocyte spike-in results in the death of 21.9% of leukocytes. Affirming the results from fig. 3a, this emphasizes the consistency and reproducibility of experimental outcome using capacitive coupling. While the standard rationale invokes the inhibition of unspecific side effects due to electrochemistry, this does not serve as an explanation for the observed specificity in this capacitively coupled system (Manuscript accepted by *Lab on a Chip*, will be cited in the final version). A further proposed mechanism for the specificity of electric field mediated lysis relates to the expansion of affected nuclei and feasibly explains the lower lysis threshold of malignant cells [10]. However, applied to our findings, these features do not reflect the lysis dynamics of erythrocytes compared to the tested leukaemia and circulating tumour cell models. These results stress the necessity of a mechanistic explanation for the dynamics of cell lysis using capacitive coupling of electric fields.

## 4. Conclusion

The current understanding of electroporation implies a selective component to the interaction of electric fields with lipid double layers. Despite this theoretical selectivity, a practical approach for electric-field mediated cell discrimination in liquid samples comprises significant challenges. While tumour ablation in tissues can be achieved in a selective manner by careful configuration of spatial electrode geometry, no comparable workaround is available for cell suspensions. Suspended cells subjected to electric fields applied with capacitive coupling have a consistent response that results in reproducible outcome with low standard deviation. This enables the selective lysis of specific cell types from a mixture of cells. Surprisingly, the order of lysis observed for the tested cell types does not coincide with pertinent literature. Similar lysis dynamics are observed for MCF-7 and Jurkat cells despite their significant size difference. These and other discussed findings are in conflict with previous reports on the subject and imply a further feature of electric field susceptibility not related to cell radius. This indicates that the potential of selective lysis by electric fields may offer greater flexibility than previously assumed. We are currently invested to reconciliate these findings by thorough characterization of the different lysis mechanics between commonly used platforms and capacitively coupled systems.

## Author contributions

Conceptualization, T.W., E.K.E. and K.J.W.; Methodology, T.W., V.K.K and K.J.W.; Investigation, T.W. and V.K.K.; Resources, K.J.W. and E.K.E.; Writing – Original Draft, T.W.; Writing – Review & Editing, T.W., E.K.W. and K.J.W.; Visualization, T.W. and V.K.K; Supervision, K.J.W. and E.K.W.; Project Administration, K.J.W; Funding Acquisition, K.J.W.

## Acknowledgement

The authors would like to acknowledge the Austrian Institute of Technology GmbH as the source of funding for this Project. Thanks go to the FFG for funding the internship of Verena K. Köhler. We acknowledge Seta Küpcü from the Department of Nanobiotechnology at BOKU University for her support in flow cytometry data acquisition.

